# Model-Driven Hybrid AI Framework for End-to-End Autonomous Decision-Making in Drug Development

**DOI:** 10.64898/2026.02.03.703154

**Authors:** Hyunjung Lee, Hyeonseok Kang, Woojin Jung, Hyojin Cho, Sungwoo Goo, Hwi-yeol Yun, Min-Gul Kim, Jung-woo Chae, Sang-Min Park, Soyoung Lee, Jae Hyun Kim, Sangkeun Jung

**Author notes:** These authors contributed equally to this work (Hyunjung Lee, Hyeonseok Kang, Woojin Jung). Co-corresponding authors: Jung-woo CHAE Sang-Min Park, Soyoung Lee, Jae Hyun Kim, Sangkeun Jung.

## Abstract

Decision-making in drug development spans heterogeneous stages from molecular design to clinical optimization, yet computer-aided workflows across stages remain fragmented, limiting traceability from evidence-derived clinical questions to simulation scenarios and decision endpoints. We present a model-driven hybrid AI framework for end-to-end decision support that treats PICO (Participants, Intervention, Comparison, Outcome) as a machine-actionable specification to define scenarios and align endpoints across modules.

Given a PICO-defined clinical question and a drug SMILES, the framework infers PBPK-ready parameters from structure, generates exposure via mechanistic PBPK simulation, and links exposure outputs to clinic-facing therapeutic drug monitoring (TDM) decision support using an open-source NLME-based Clinical Pharmacokinetic Consultant Service (CPCS). We additionally implement an LLM-based PICO extraction layer to structure clinical abstracts into P/I/C/O elements for systematic evidence ingestion.

Results include backbone verification for DTI-guided intrinsic clearance prediction under fold-error bounds and PBPK trajectory plausibility, and external validation of the hybrid AI–PBPK pipeline showing substantial dispersion for several endpoints. An input-controlled comparison between PhysioSim and PK-Sim on *N* = 9 drugs yields similarly degraded accuracy under identical AI-predicted inputs, suggesting upstream parameter quality as a dominant constraint. For evidence ingestion, prompting-only PICO extraction achieves F1 of 0.687 on EBM-NLP and 0.552 on TB-PICO. For clinical utility, CPCS improves TDM performance over a PKS baseline on phenobarbital and vancomycin, reducing MAPE by approximately 26–60% depending on configuration.

Overall, the framework provides a modular, traceable blueprint linking evidence-defined questions to exposure simulation and TDM decision support while preserving mechanistic interpretability. Future work will integrate outcome-aligned time-to-event modeling and federated learning to close the PICO-to-outcome loop and improve parameter inference under multi-site governance.

## 1 Introduction

### 1.1 Challenges in end-to-end decision-making across the drug development lifecycle

Drug development is an inherently complex, multi-stage process spanning molecular design, preclinical evaluation, clinical development, and regulatory assessment, with high cost and low overall success rates reported across clinical phases [1, 2]. Across these stages, decision-making is often fragmented due to heterogeneous data sources, stage-specific modeling paradigms, and sequential workflows that rely heavily on manual integration and expert heuristics. Despite substantial advances in computational modeling, a persistent bottleneck remains in pharmacokinetics translation: exposure prediction often requires extensive experimental absorption, distribution, metabolism, and excretion (ADME) characterization and iterative model refinement, which constrains throughput and hinders systematic exploration of design and dosing strategies [3].

### 1.2 Limitations of data-driven and mechanistic approaches

Recent AI-driven approaches have demonstrated notable performance in isolated tasks such as property prediction and outcome modeling. However, purely data-driven models often lack mechanistic interpretability and struggle to generalize beyond the domains represented in training datasets, which limits reliability in high-stakes settings [4]. Conversely, model-informed approaches grounded in pharmacokinetics and systems pharmacology provide mechanistic coherence and regulatory credibility, but are frequently constrained by high dimensionality, parameterization burden, and practical challenges in decision contexts [5, 6]. As a consequence, neither data-driven nor mechanistic approaches alone are sufficient to support scalable and autonomous decision-making across the full drug development pipeline.

In evidence-based pharmacotherapy, systematic reviews and meta-analyses commonly define clinical questions using structured PICO elements (Participants, Intervention, Comparison, Outcome), which specify the target population, comparator regimen, and clinically meaningful endpoints for decision-making [7, 8]. However, model-driven pipelines—such as physiologically based pharmacokinetic (PBPK) exposure simulation, therapeutic drug monitoring (TDM) support, and outcome-oriented modeling—often optimize surrogate PK endpoints (e.g., AUC/*C*_max_) without explicitly importing the PICO specification that grounded the evidence synthesis. This can reduce end-to-end traceability from published evidence to simulation scenarios and event definitions in outcome models.

To address this gap, we treat PICO not merely as a literature-search aid, but as a *machine-actionable specification* that governs (i) scenario instantiation for simulation and comparison (P/I/C) and (ii) endpoint alignment for downstream decision-making (O) [7, 8]. Under this view, PICO appears twice in an end-to-end loop: it initiates model-based analysis by defining the clinical question and comparison scenarios, and it re-enters at the final stage by anchoring outcome-oriented decision targets (planned via time-to-event modeling). This perspective motivates placing PICO at the top of the methodological stack rather than as an auxiliary post-processing step.

### 1.3 Rationale for a model-driven hybrid AI framework

This work targets a practical middle ground: a hybrid approach that preserves mechanistic structure while minimizing experimental dependency. We propose a modular pipeline: **SMILES** →**AI-based ADME/clearance inference**→ **mechanistic PBPK simulation** →**clinical decision modules** →**planned TTE**. To address complementary limitations of learning-based and mechanistic approaches, the framework integrates AI components as parameter generators and PBPK as a physiologically grounded decision backbone [9, 5].

Accordingly, we design the framework so that evidence-derived **PICO** specifications define the simulation and comparison scenarios upfront, while the resulting exposure outputs are carried forward to TDM decision support and (in future work) re-mapped to outcome targets via TTE modeling.

### 1.4 Contributions

Our contributions are as follows:

- We propose an end-to-end traceable framework that positions PICO as a top-level specification linking evidence-derived questions to model scenarios and decision endpoints.
- We implement a molecular-to-exposure pipeline that combines SMILES-driven parameter inference with an ODE-based PBPK simulator to generate clinically interpretable exposure profiles.
- We benchmark PBPK behavior through distribution model comparisons and report external validation of the hybrid pipeline on an independent drug set.
- We demonstrate downstream clinical utility via Clinical Pharmacokinetic Consultant Service (CPCS) benchmarks against a widely used TDM baseline (PKS) and outline a planned time-to-event module to align decisions with PICO Outcomes.

### 1.5 Objectives and scope of the study

The goal is to establish an end-to-end molecular-to-exposure module that: (i) predicts ADME properties from SMILES, (ii) predicts metabolism-enzyme interactions and intrinsic clearance, (iii) integrates these parameters into PBPK to simulate clinically interpretable endpoints (e.g., *C*_max_, AUC, and *T*_max_), and (iv) connects exposure outputs to a clinic-facing TDM decision module (CPCS). TTE integration is described as planned future work.

Figure 1 provides an end-to-end overview of the proposed workflow, highlighting how PICO-guided scenario specification is connected to Hybrid AI–PBPK exposure simulation and CPCS-based clinical decision support, with time-to-event modeling planned to align outcomes with the original PICO definition.

**Figure 1:**
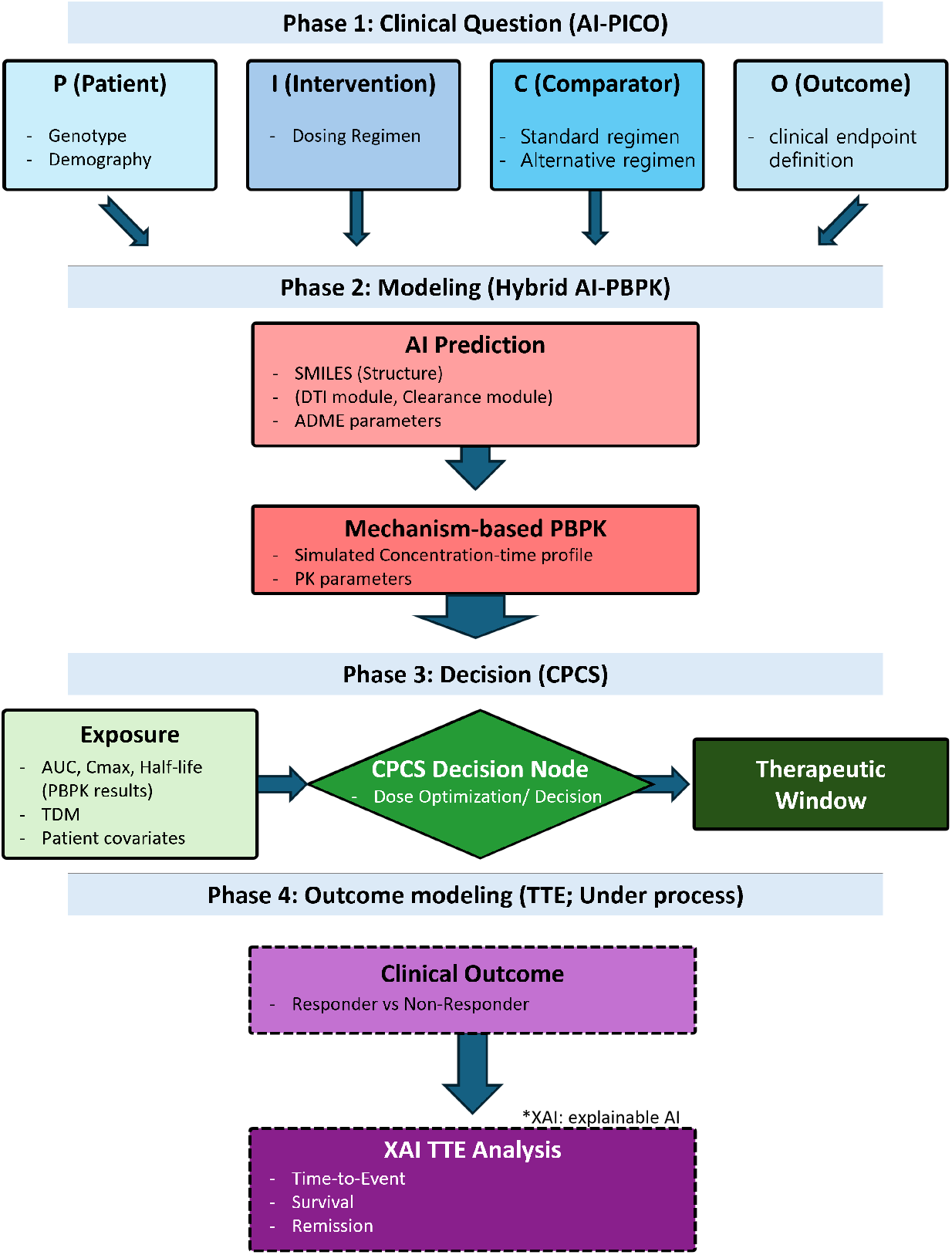
End-to-end framework for traceable decision-making in drug development. AI–PICO structures clinical questions, Hybrid AI–PBPK translates them to exposure, and CPCS supports dose decisions; outcome modeling (TTE) is under development.

## 2 Methods

### 2.1 Overall architecture of the model-driven hybrid AI framework

The framework consists of four modules connected by explicit interfaces. Inputs are (i) a PICO-defined clinical question (Participants, Intervention, Comparison, Outcome) that specifies the target population, dosing/comparator scenarios, and endpoint targets, and (ii) drug chemical structure represented as SMILES. Outputs are standardized exposure summaries and concentration forecasts used for TDM decision support, with a planned time-to-event module that maps exposure and covariates to outcome-oriented event risks. Across modules, we pass standardized objects: SMILES and derived descriptors; PBPK-ready parameters; simulated concentration–time profiles and exposure summaries; and patient-level covariates and TDM observations. Figure 1 summarizes the end-to-end workflow, while Figure 2 details the module interfaces and data flow across evidence ingestion, Hybrid AI–PBPK modeling, and CPCS-based decision support (with TTE planned as an extension).

**Figure 2:**
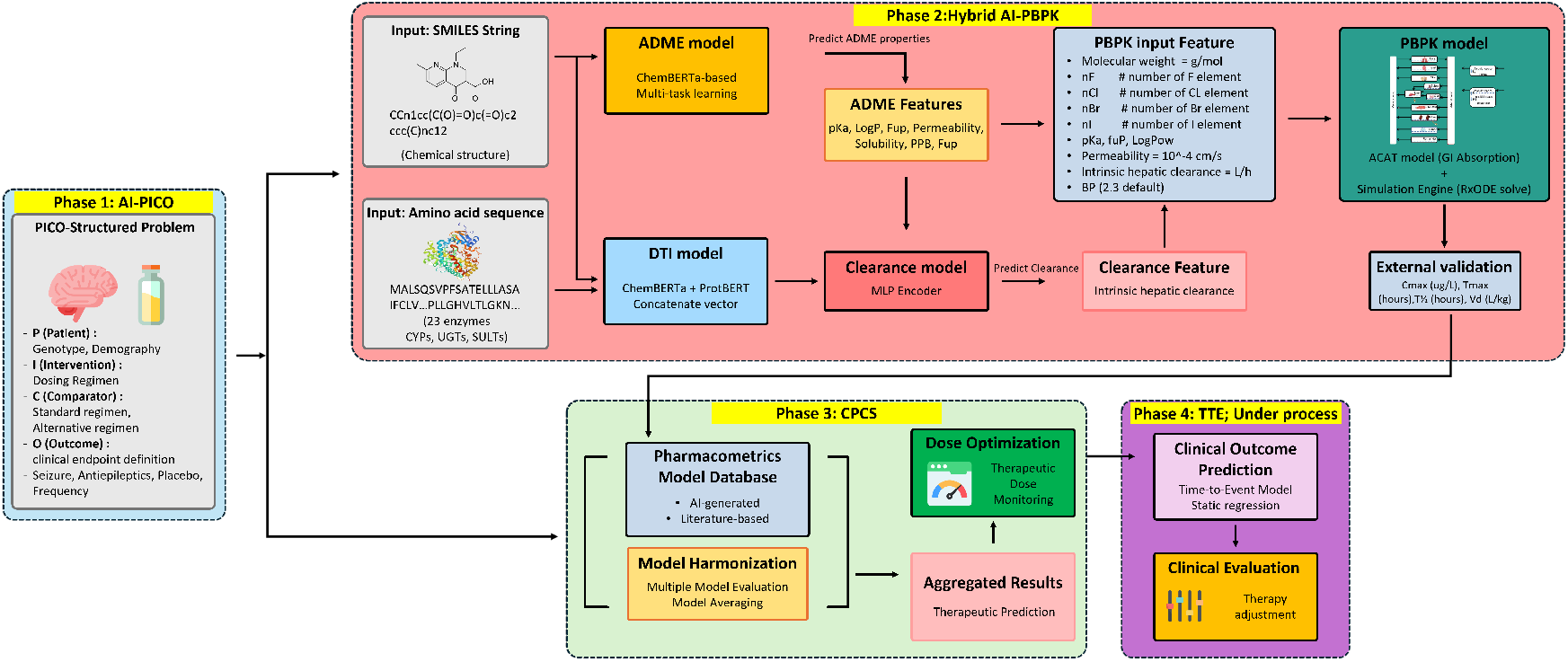
Detailed Hybrid AI–PBPK pipeline used in the framework. Molecular inputs (e.g., SMILES/sequence) are converted to ADME/clearance features for PBPK simulation and external PK validation.

### 2.2 Evidence ingestion and PICO-guided decision specification

We use PICO as a machine-actionable specification to ensure traceability from evidence-derived questions to simulation scenarios and decision endpoints. Participants define cohort constraints and covariates (e.g., renal function, age bands, pregnancy status); Intervention defines drug and dosing regimen; Comparison defines alternative regimen(s) or standard-of-care settings; and Outcome defines clinically meaningful targets that will be mapped to event definitions in the planned time-to-event module. This specification is used to parameterize simulation setups (e.g., dosing schedules, population settings) and to declare the endpoint alignment between exposure-based surrogates (e.g., AUC/*C*_max_) and outcome-oriented targets.

To operationalize PICO from unstructured clinical text, we implement an LLM-based extraction procedure that converts abstracts and clinical summaries into structured Participants/Intervention/Comparison/Outcome elements [8, 7, 10]. We adopt prompting without task-specific fine-tuning [10, 11], incorporating similarity-based few-shot selection [12, 13] and brief prompt enrichment [14]. We denote this configuration as SS+EB, where SS indicates similarity-selected few-shot prompting and EB indicates biomedical prompt enrichment. Extraction performance is evaluated on EBM-NLP [15] and TB-PICO, reporting F1 aggregated over PICO elements and comparing against representative fine-tuned baselines [16, 17, 18].

### 2.3 SMILES-to-exposure modeling: parameter prediction and PBPK simulation

Figure 2 outlines the SMILES-to-parameter prediction stack and its interface to the ODE-based PBPK simulator. Heterogeneous ADME datasets totaling 39,115 unique compounds were aggregated from EU-KIST, the Drug–Drug Interaction Database (DIDB) [19], and the K-MELLODDY consortium. All compound identifiers were standardized at the SMILES level, and experimental records were harmonized through consistent units and quality control. Regulatory-facing discussions on artificial intelligence for drug development informed the motivation for learning-enabled components in the workflow [20]. Key ADME-related properties are inferred from SMILES using an ML model trained on multi-source compound collections.

#### SMILES-based ADME parameter inference

We predict PBPK-relevant ADME parameters directly from SMILES to reduce dependence on experimental characterization. SMILES strings are encoded with a transformer-based molecular language model, and the resulting representation is fed to a multi-task regression head that outputs key inputs required by the PBPK simulator. Predicted properties include lipophilicity (LogP), ionization (p*K*_*a*_), aqueous solubility, permeability, plasma protein binding, and in-vitro fraction unbound (*f*_*u*_). Because real-world ADME datasets are sparsely labeled across endpoints, we use endpoint-wise masking so that each compound contributes loss only for properties with available ground truth, enabling joint training without discarding partially labeled records. These SMILES-derived ADME estimates are then mapped to PBPK parameters governing absorption (e.g., solubility/permeability-limited uptake), distribution (e.g., protein binding and tissue partitioning), and systemic exposure summaries.

Predicted properties include physicochemical and biopharmaceutic features (e.g., partitioning, permeability, solubility, and fraction unbound in plasma) and are used as PBPK inputs under limited experimental availability.

#### Mechanism-aware metabolism and clearance inference

To reduce the parameterization burden for PBPK elimination terms, we model metabolism in two coupled steps: (i) drug–enzyme interaction inference and (ii) intrinsic clearance regression conditioned on predicted enzyme involvement, the clearance predictor estimates intrinsic clearance *CL*_*int*_ (in log-space for stability), which is subsequently mapped to PBPK elimination parameters via standard hepatic clearance assumptions.

#### Mechanistic PBPK simulation

PBPK is implemented as an ODE system and solved numerically using rxode2 in R [21, 22]. The simulator comprises organ compartments with blood-flow coupling and tissue distribution, and absorption is represented with a GI component including solubility-related effects. Primary outputs include plasma concentration–time curves and derived endpoints such as AUC and *C*_max_ (and when feasible *T*_max_, *V*_*d*_, and half-life) [9, 5]. Evaluation is conducted on independent compounds using mean squared error (MSE), root mean squared error (RMSE), *R*^2^, mean absolute error (MAE), Pearson correlation, and fold-error coverage (within 2-fold and 3-fold).

### 2.4 Integrated clinical decision optimization and planned outcome prediction

Clinical decision support is implemented as CPCS, an open-source TDM platform built on a nonlinear mixed-effects (NLME) engine that supports PBPK, and published PK (and when available PK/PD) models with covariates. CPCS operates on longitudinal patient records (dosing history, concentrations, covariates) and performs repeated estimation–forecast cycles as new TDM measurements are incorporated. We benchmark CPCS against a widely used TDM software standard PKS [23] on phenobarbital and vancomycin models and datasets [24, 25], reporting MAPE/MPE and F20/F30 under repeated updates, along with qualitative robustness observations (e.g., numerical failures and runtime burden).

## 3 Results

### 3.1 PICO extraction performance

We evaluate the proposed LLM-based PICO extraction layer as an evidence-ingestion component that converts unstructured abstracts into structured Participants/Intervention/Comparison/Outcome elements. Because downstream modules (scenario instantiation for PBPK and endpoint alignment for planned outcome modeling) depend on correct PICO specification, we report extraction accuracy using span-level F1 aggregated over PICO elements.

Table 1 summarizes results on EBM-NLP and TB-PICO. On EBM-NLP, the proposed prompting strategy (SS+EB) improves over zero-shot and random few-shot prompting, reaching F1 = 0.687 and narrowing the gap to a fine-tuned BioLinkBERT baseline (F1 = 0.723). On TB-PICO, the same strategy yields F1 = 0.552, exceeding zero-shot performance and improving over random few-shot prompting. Overall, SS+EB prompting consistently improves extraction F1 over zero-shot and random few-shot settings across both datasets, reducing the gap to a fine-tuned BioLinkBERT baseline on EBM-NLP and yielding the best prompting-only performance on TB-PICO.

**Table 1:** PICO extraction performance (F1) comparing prompting methods and fine-tuned baseline models.

| Dataset | Model/Method | Setting | F1 |
| --- | --- | --- | --- |
| EBM-NLP | BioLinkBERT-large | Fine-tuned | 0.723 |
|  | ChatGPT | Zero-shot | 0.583 |
|  | ChatGPT | Few-shot (Random) | 0.626 |
|  | ChatGPT | Proposed (SS+EB) | 0.687 |
| TB-PICO | BioLinkBERT-large | Fine-tuned (EBM-NLP+TB-PICO) | 0.494 |
|  | ChatGPT | Zero-shot | 0.438 |
|  | ChatGPT | Few-shot (Random) | 0.489 |
|  | ChatGPT | Proposed (SS+EB) | 0.552 |

### 3.2 SMILES-to-ADME parameter prediction

To confirm that SMILES-based inference provides PBPK-ready inputs, we evaluated the ADME parameter predictor on held-out compounds. Figure 3 summarizes SMILES-to-ADME predictions for PBPK inputs across six endpoints: LogP, p*K*_*a*_, aqueous solubility, membrane permeability, plasma protein binding (PPB), and in-vitro fraction unbound (*f*_*u*_). To improve comparability across heterogeneous sources, several endpoints are evaluated on a log-transformed scale (as indicated on the figure axes), so the scatter should be interpreted in log space rather than raw units. These predicted ADME properties are subsequently mapped to PBPK-ready parameters governing absorption, distribution, and binding in the downstream simulator. Overall performance varied by endpoint, with *R*^2^ ranging from 0.177 to 0.810, RMSE from 0.277 to 0.894, and Pearson *r* from 0.446 to 0.900; accuracy was strongest for LogP (*R*^2^ = 0.810, *r* = 0.900) and moderate for solubility/permeability, while p*K*_*a*_ showed lower agreement (*R*^2^ = 0.177, *r* = 0.446).

**Figure 3:**
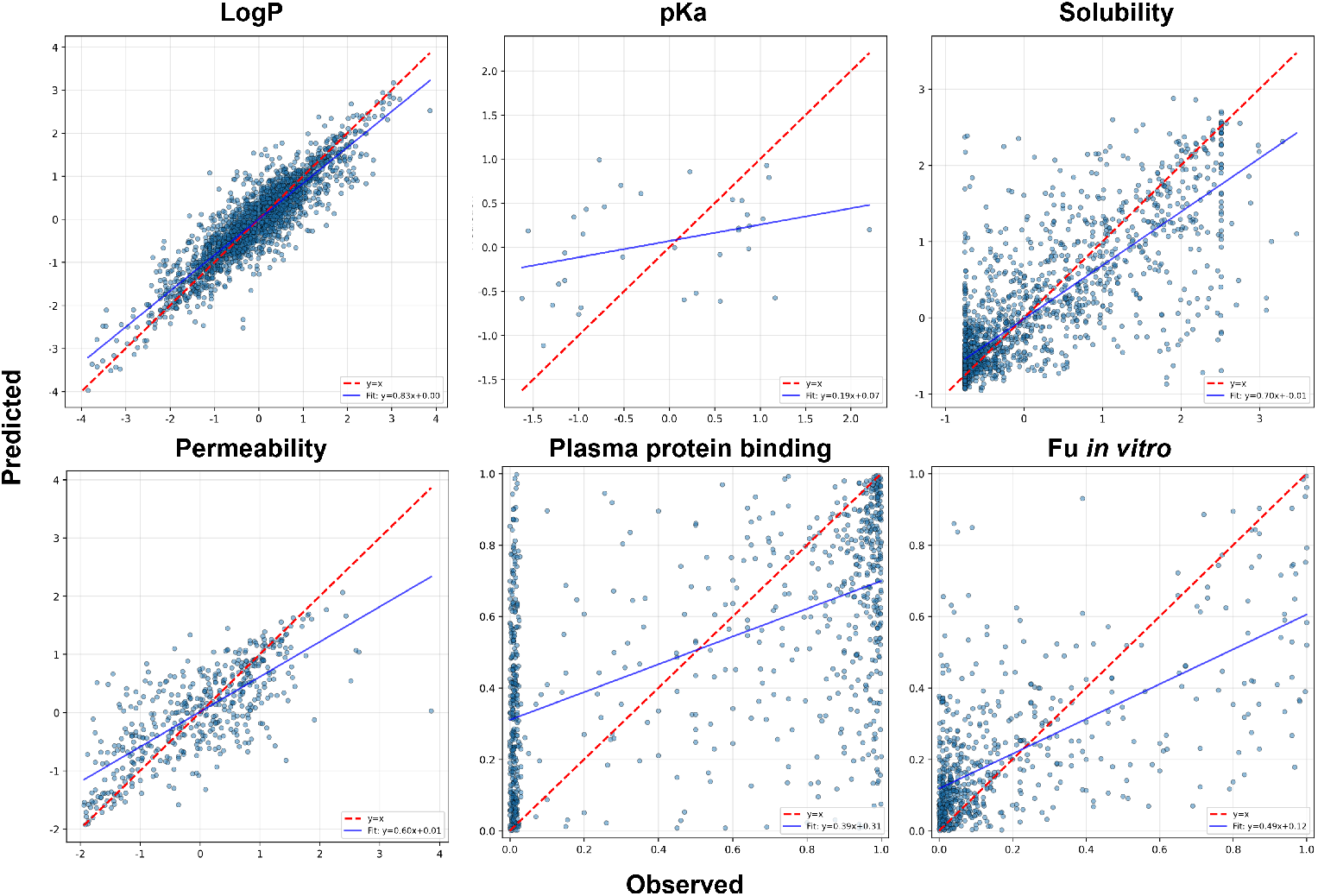
SMILES-to-ADME parameter prediction on a held-out test set. Scatter plots show predicted versus observed values across multiple ADME properties used as PBPK inputs. The solid line indicates unity (*y* = *x*). Several endpoints are shown on a log-transformed scale as indicated on the axes.

### 3.3 Backbone verification for clearance inference and PBPK simulation

Before analyzing end-to-end AI–PBPK performance, we first validate the upstream clearance-related component and then confirm time-domain plausibility of the PBPK backbone. Figure 4 summarizes these backbone-level checks.

**Figure 4:**
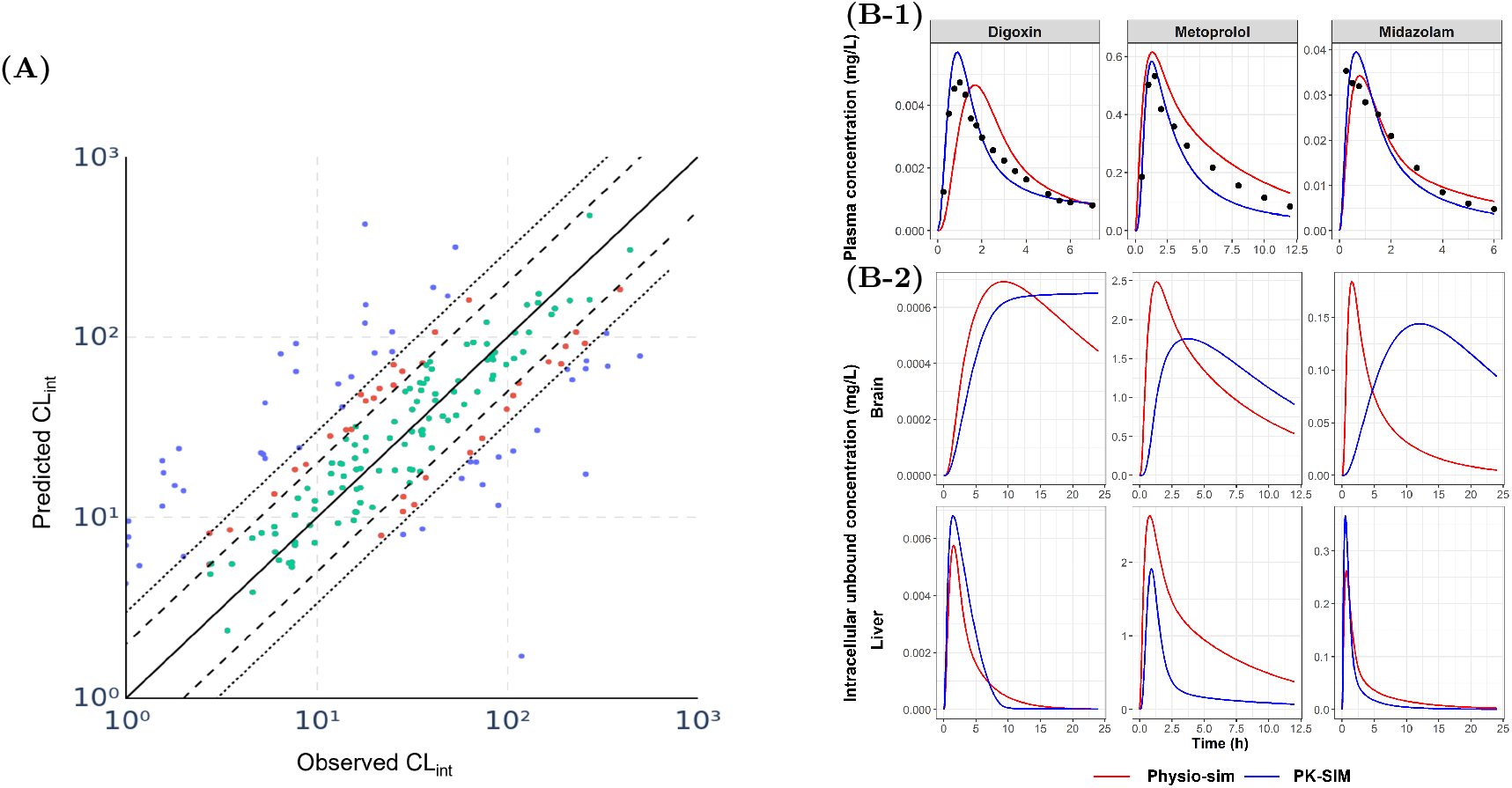
Backbone verification of clearance inference and PBPK simulation. (A) Predicted vs. observed *CL*_*int*_ on a log–log scale; the solid line indicates unity (*y* = *x*), and dashed/dotted lines indicate 2-fold and 3-fold error bounds. (B-1) Plasma concentration–time profiles for representative compounds; points indicate observed measurements and lines indicate model predictions (Physio-Sim vs. PK-Sim).(B-2) Simulated tissue concentration–time profiles without observed points: brain (upper row) and liver (lower row), shown as intracellular unbound concentrations over time for the same compounds.

#### DTI-guided intrinsic clearance prediction under fold-error bounds

Figure 4(A) evaluates intrinsic clearance prediction under fold-error bounds. We report accuracy using the fold ratio *predicted CL*_*int*_*/observed CL*_*int*_ and show that a substantial fraction of compounds falls within 2-fold and 3-fold regions, while remaining deviations are dominated by outliers that may reflect route misspecification, sparse enzyme annotations, or unmodeled non-hepatic pathways. As a reference baseline, the clearance predictor achieves RMSE = 2.413 with 52.97% and 70.79% of compounds within 2-fold and 3-fold error, respectively (*N* = 172).

#### Verification of the PBPK simulation backbone in the time domain

We next provide a sanity check to confirm that the PBPK backbone produces plausible concentration–time trajectories when driven by the same upstream AI-derived inputs. Figure 4(B-1) shows representative plasma profiles with observed clinical concentrations (points) and model predictions (lines), comparing PhysioSim and PK-Sim under identical inputs. In addition, Figure 4(B-2) reports tissue simulations (brain and liver) for the same compounds to illustrate mechanistic plausibility of distribution at the organ level; these tissue trajectories are shown without observed points because matched tissue measurements are not available in this evaluation.

### 3.4 External validation of the hybrid AI–PBPK pipeline

We evaluate the end-to-end hybrid AI–PBPK pipeline on an independent drug set. Figure 5(A) reports parity plots under fold-error bounds, indicating substantial dispersion for several endpoints despite complete profile generation from AI-predicted inputs. To test whether the observed errors are driven primarily by the PBPK engine or by AI-derived PBPK input parameters, we perform an input-controlled comparison: we inject the same AI-predicted PBPK inputs into two PBPK engines—the PhysioSim backbone used in this study and the commercial PK-Sim platform—and compute endpoint-wise performance.

**Figure 5:**
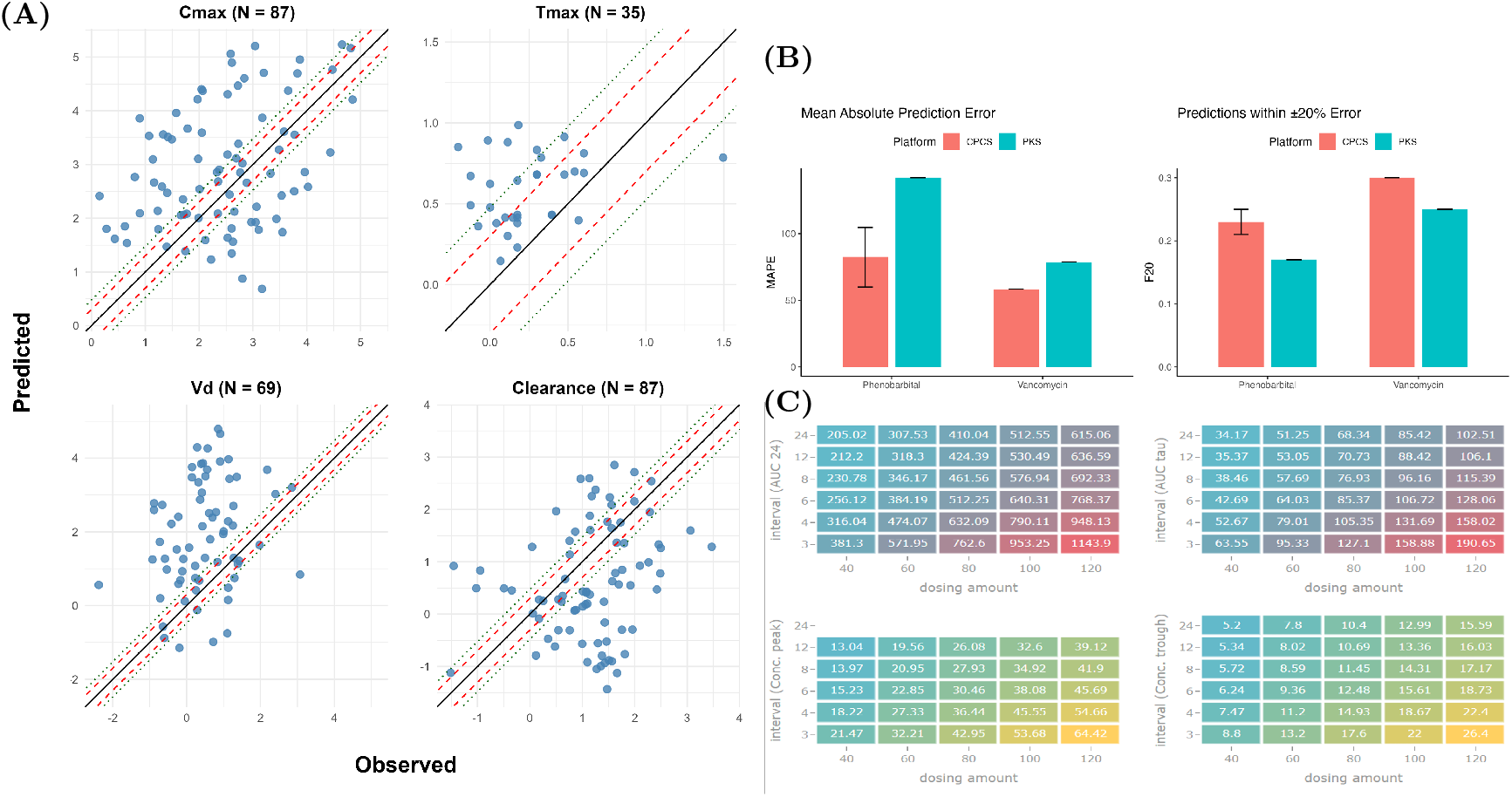
End-to-end evaluation: external validation of Hybrid AI–PBPK and downstream TDM decision support. (A) External validation parity plots across endpoints; the solid line indicates unity (*y* = *x*), dashed lines indicate 2-fold error bounds, and dotted lines indicate 3-fold error bounds. (B) CPCS benchmark summary against the PKS baseline using MAPE and F20 across drugs. (C) Example heatmap illustrating dose selection/optimization across dosing amounts and dosing intervals.

As shown in Table 2, both engines exhibit similarly degraded accuracy under identical AI-derived inputs for multiple endpoints, suggesting that upstream parameter quality and model-mismatch (e.g., distribution and elimination assumptions) are dominant constraints.

**Table 2:** PhysioSim vs. PK-Sim endpoint performance using identical AI-predicted PBPK inputs. The PK-Sim re-implementation required substantial manual model construction; therefore, the comparison was conducted on *N* =9 randomly selected drugs (and *N* =4 for *T*_max_ where available).

| Model | Parameter | $N$ | MSE | RMSE | $R^2$ | MAE | Pearson $r$ | %2-fold | %3-fold |
| --- | --- | --- | --- | --- | --- | --- | --- | --- | --- |
| PhysioSim | $C_{\max}$ ($\mu\text{g/L}$) | 9 | 1.413 | 1.189 | 0.355 | 0.981 | 0.622 | 22.2 | 22.2 |
| | $T_{\max}$ (h) | 4 | 0.103 | 0.321 | -1.168 | 0.301 | 0.861 | 50.0 | 100.0 |
| | $V_d$ (L/kg) | 9 | 3.583 | 1.893 | -22.248 | 1.618 | 0.161 | 11.1 | 22.2 |
|  | Clearance (L/h) | 9 | 1.783 | 1.335 | -3.000 | 0.847 | -0.139 | 44.4 | 55.6 |
| PK-Sim | $C_{\max}$ ($\mu\text{g/L}$) | 9 | 9.458 | 3.075 | -2.211 | 2.711 | 0.097 | 0.0 | 11.1 |
| | $T_{\max}$ (h) | 4 | 0.037 | 0.193 | -15.412 | 0.173 | -0.826 | 75.0 | 100.0 |
| | $V_d$ (L/kg) | 9 | 22.447 | 4.738 | -7.233 | 4.310 | 0.175 | 0.0 | 0.0 |
|  | Clearance (L/h) | 9 | 6.046 | 2.459 | -3.703 | 1.962 | 0.077 | 11.1 | 11.1 |

### 3.5 CPCS benchmarks for therapeutic drug monitoring decision support

We benchmark CPCS as a clinic-facing concentration prediction and decision-support component under therapeutic drug monitoring (TDM), where predictions are repeatedly updated as new concentration measurements become available. We compare CPCS against a widely used TDM software baseline (PKS standard) on phenobarbital and vancomycin datasets, reporting mean absolute prediction error (MAPE) and threshold accuracy within ±20% relative error (F20). Figure 5(B–C) summarizes the CPCS benchmarking results and an example dose-selection scenario.

## 4 Discussion

This work demonstrates a traceable blueprint from chemical structure to clinically meaningful outputs, with PICO anchoring both scenario specification and endpoint alignment. The hybrid AI–PBPK design preserves mechanistic interpretability by keeping physiological structure explicit while using ML to supply hard-to-measure parameters from SMILES and enzyme-related inputs [9, 5]. In parallel, the PICO layer bridges clinical literature and decision logic, supporting auditability when translating evidence into dosing and outcome strategies. From a system perspective, these results suggest that a prompting-only PICO extractor can serve as a lightweight evidence-structuring layer without task-specific fine-tuning. For downstream clinical utility, CPCS shows improved TDM performance over a PKS baseline on phenobarbital and vancomycin, as reflected by lower MAPE and higher F20. Qualitative observations suggest CPCS is more robust under repeated TDM updates, where PKS may exhibit prediction failures or heavy computational burden. External validation indicates limited fold-error coverage for several endpoints, reflecting error propagation from predicted solubility/unbound fraction and incomplete representation of non-hepatic elimination pathways (e.g., renal clearance). Improving upstream property prediction, adding multi-route elimination, and expanding dataset diversity are expected to be high-impact steps. While the framework is designed end-to-end, the current evaluation demonstrates feasibility of (SMILES → AI–PBPK) and separate benchmarking of downstream modules (CPCS, PICO), rather than a fully integrated exposure-to-outcome evaluation. Two extensions are planned to complete the end-to-end traceability loop and to improve sustainability of the learning components. First, a time-to-event (TTE) module will re-connect exposure-based surrogates to PICO Outcomes by mapping PBPK- and/or TDM-updated exposure summaries and covariates to individualized hazards and survival probabilities under candidate dosing strategies. Second, we plan to incorporate federated learning for the SMILES-to-parameter modules to support multi-site model evolution without centralizing sensitive data.

## 5 Conclusion

We presented a model-driven hybrid AI framework that connects SMILES-driven inference, mechanistic PBPK simulation, and clinic-facing TDM decision support (CPCS), with a planned TTE module to re-align final decisions with PICO Outcomes. Results show backbone verification (DTI-guided clearance prediction under fold bounds and PBPK trajectory plausibility), limitations observed in hybrid AI–PBPK external validation, strong PICO extraction performance without fine-tuning, and CPCS benchmarks demonstrating improvements over a widely used TDM baseline. This modular design provides a transparent and extensible foundation for end-to-end, explainable decision support across drug development stages. Future work will integrate an outcome-aligned TTE module and federated learning to complete the PICO-to-outcome traceability loop and support sustainable multi-site model refinement.

